# Genome Elimination Mediated by Gene Expression from a Selfish Chromosome

**DOI:** 10.1101/793273

**Authors:** Elena Dalla Benetta, Igor Antoshechkin, Ting Yang, Hoa Quang My Nguyen, Patrick M. Ferree, Omar S. Akbari

## Abstract

Numerous plants and animals harbor selfish B chromosomes that “drive,” or transmit themselves at higher-than-Mendelian frequencies, despite long-term fitness costs to the organism. Currently it is unknown how B chromosome drive is mediated, and whether B-gene expression plays a role. We used modern sequencing technologies to analyze the fine-scale sequence composition and expression of Paternal Sex Ratio (PSR), a B chromosome in the jewel wasp *Nasonia vitripennis*. PSR causes female-to-male conversion by destroying the sperm’s hereditary material in young embryos in order to drive. Using RNA interference, we show that testis-specific expression of a PSR-linked gene, named *haploidizer*, facilitates this genome elimination-and-sex conversion effect. *Haploidizer* shares homology with a gene in *Candidatus cardinium*, a bacterial symbiont that also induces genome elimination in its insect host.

**One Sentence Summary:** *haploidizer* mediates B chromosome drive

## Main Text

In eukaryotes, the genome contains a core set of chromosomes that are indispensable for viability. However, thousands of plant and animal species also harbor non-essential chromosomes termed B chromosomes, consisting of both coding and non-coding sequences thought to be mostly copied from the essential, or A, chromosomes. Despite their non-essential nature, B chromosomes presumably carry no needed genes and therefore are, in principle, prone to loss over successive generations (*1*–*5*). To counter the tendency to be lost, many B chromosomes possess the extraordinary ability to “drive,” or transmit themselves at frequencies above those predicted by Mendelian rules (reviewed in (*6, 7*)). Although drive is necessary for B chromosome transmission, it can impose deleterious effects on the inheritance of other genomic regions, thus having potentially strong influences on organismal fitness (*5*, *6*, *8*). Currently, it is not understood how B chromosomes can behave so differently in their transmission from other, non-driving chromosomes.

One of the most striking examples of drive is caused by a B chromosome known as PSR (for Paternal Sex Ratio), which has been detected at moderate frequencies (6-11%) within certain populations of the jewel wasp *Nasonia vitripennis (9)*. PSR is transmitted strictly paternally (*i.e*., via sperm), and its presence leads to the complete elimination of the sperm’s essential chromosomes, but not PSR itself, during the mitotic division immediately following fertilization (*10*). Genome elimination results directly from failure of the sperm’s chromatin to resolve into individualized chromosomes upon entry into mitosis and is preceded by distinct chemical markers that indicate a disrupted chromatin state (*11*–*14*). This effect is critical for the transmission of PSR. Wasps, like all hymenopteran insects (also including ants and bees), reproduce through haplo-diploidy, in which unfertilized eggs develop into haploid males while fertilized eggs become diploid females. Therefore, elimination of the sperm’s hereditary material by PSR converts all diploid, female-destined eggs into haploid males, the PSR-transmitting sex (*9*). Broadly, this effect results in severely male-biased sex ratios at the deme level, which can negatively impact wasp metapopulations (*15*).

A fundamental biological question is how PSR and, more broadly, other B chromosomes are capable of interacting uniquely with the cellular environment in order to drive. Previous studies identified three distinct ~180 DNA-base pair repeats, termed PSR2, PSR18, and PSR22, which are abundant in copy number on PSR, but not present on any of the five essential wasp chromosomes (*16*, *17*). Each of these repeats contains a highly conserved, 8-bp palindromic motif that is reminiscent of those found in some transcription factor-binding sequences (*16*, *17*). These characteristics have led to speculation that PSR-specific sequences may act as a sink for some limited chromatin factor(s), drawing them away from the rest of the genome and causing defective chromatin structure (*16*). An underlying aspect of this scenario is that the driving effect of PSR is the result of intrinsic properties that are unique to its sequence composition. For this reason, we refer to such an effect as *passive*. Alternatively, PSR’s drive may involve the *active* expression of B-linked sequences. Recent studies have begun to identify individual sequences that are expressed from a number of different B chromosomes, including PSR (*18*, *19*). While no studies have, to date, demonstrated functionality for any B-linked sequence, it is plausible that an RNA or protein expressed by PSR functions as an effector of drive by disrupting normal transmission of the paternal chromatin.

Previous transcriptional profiling uncovered a handful of long, polyadenylated RNAs expressed uniquely by PSR in the wasp testis (*18*). To further this work, we used modern genome sequencing technologies to generate large scaffolds of the PSR chromosome, onto which we mapped these PSR-expressed transcripts. These efforts produced a comprehensive portrayal of PSR’s sequence composition, models for its individual genes, and its transcribed loci. Using RNA interference (RNAi), we show that genome elimination requires the expression of one of these PSR-linked loci, demonstrating an active (*i.e.*, gene expression-based) involvement of this B chromosome in its own drive.

To deduce PSR’s sequence composition, we sequenced the genome of both wildtype and PSR-carrying wasps separately using a combination of PacBio, Oxford Nanopore and Illumina technologies, generating a 297Mb assembly comprising 444 contigs with an N50 length of 6.6 Mb representing a 13.9 fold decrease in contig number and 9.3 fold increase in N50 length compared to the current *N. vitripennis* genome assembly (*20*). Using the hymenoptera-specific set of universal single-copy orthologs, the genome completeness is estimated to be 96.5% (Table S1) (*21*). PSR-specific contigs were identified using the chromosome quotient method, which calculates the ratio of the number of wild type and PSR+ reads mapping to a contig (Table S2, Fig. S1) (*22*)) (see Methods). This approach yielded 120 contigs (9.2 MB in total, N50 of 124 kb) that were specific to PSR (Table S3). In addition, 73 contigs totaling 272 Mb (91.58% of total genome) were assigned to the five essential *N. vitripennis* chromosomes using the previously developed genetic markers (*23*) (Fig. S2, Table S4, Data S1.tar).

To better understand the sequence composition and organization of PSR (Table S5), we performed a detailed computational assessment of PSR’s repetitive content with RepeatModeler and RepeatMasker (*24*, *25*). These analyses revealed that 89.80% of PSR is composed of repetitive DNAs (Fig.1A, Table S6). The most abundant repeats (70.32%) are complex satellites that belong to four main families. Three of these satellites, PSR2 (49.54%), PSR18 (42.64%) and PSR22 (17.22%), are specific to PSR. PSR2 and PSR18 are typically found together on gene-coding contigs and they mostly overlap, whereas PSR22 is found on different sets of contigs without coding genes. A fourth repeat, NV79 (2.71%), is located on four PSR scaffolds, not overlapping with other repeats, and also on all five essential chromosomes (Fig. 1B). Additionally, PSR contains DNA and RNA (retro-) transposable elements (TEs) (13.97%), simple repeats (0.36%) and other low complexity regions (0.03%), and uncharacterized sequences (3.81%) (Fig.1A; Table S5, S6). Two PSR contigs contain telomeric sequences that cover 3.5-3.9 kb at one of the ends of each contig. Interestingly, the telomeric repeat found on PSR and at the ends of all five A-chromosomes of *N. vitripennis* is ‘TTATTGGG,’ which is different from the canonical sequence ‘TTAGGG’ (*26*) that is found in other insect species (Fig S3).

**Fig. 1.**
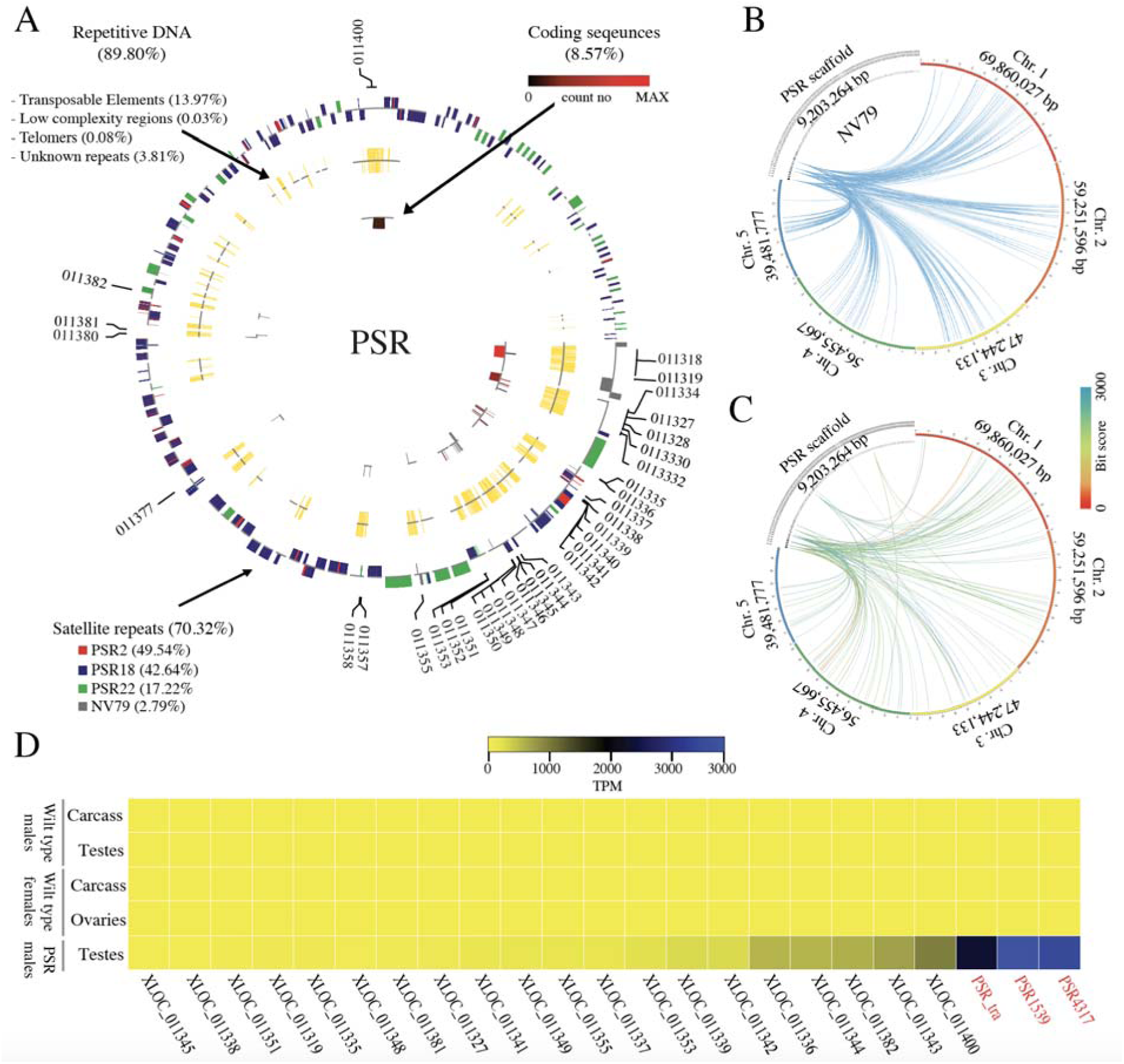
Composition and origin of PSR chromosome. **A)** Circular visualization of the PSR chromosome scaffolds. The outermost circle represents repetitive satellites with outer bars representing the positive DNA strand and the inner bars representing those on the negative DNA strand. The colors represent the four major (70.32%) satellite families. The middle track represents other repetitive sequences (17.89%) including TE’s, low complexity regions and telomeric sequences. The innermost track represents protein coding sequences (8.57%) and the color ranges from black to red for low to high expression, respectively. **B)** The relationship of NV79 repeats between PSR scaffolds and host chromosomes 1-5 indicating that PSR repeats are homologous to sequences found on all chromosomes. **C)** The relationship of PSR protein coding sequences and host chromosomes. The links are colored by blast bit scores from red (highest) and blue (lowest). Note: the PSR scaffolds are not placed in order and there is no correlation between the relationship in B and C despite the appearance of the graph. **D)** Heatmap of PSR gene with expression values higher than 10 TPM. Expression ranges from yellow to blue from low to high expression. PSR genes that were disrupted with RNAi in this study are shown in red.

Using Nanopore RNA sequencing from PSR-carrying testes and whole animals, we identified 68 transcripts (Table S7) encoded by 44 loci located on 14 of the 120 scaffolds and covering 8.57% of the total length of PSR (Table S8). The majority of the intronic regions are represented by TEs and low complexity regions; thus, the coding sequences corresponding to exonic regions represent only 0.51% of the entire PSR chromosome. Fifty of the PSR-encoded transcripts have identifiable open reading frames longer than 100 aa and 34 of those possess Pfam domains suggesting putative molecular functions that may be performed by these genes (Table S7). Fifty-eight transcripts produced significant blast hits when searched against the non-redundant protein database. Thirty-one sequences had matches to proteins encoded by the wild-type *N. vitripennis* genome, although *N. vitripennis* genes were the best hits for only 13 of them. The corresponding genes are located in different regions across all 5 essential chromosomes (Fig. 1C) and span a range of functional groups, including transposon activity, transcription regulation, DNA binding, and protein binding. Importantly, 20 transcript sequences were best matched to genes found in the genome of a closely related parasitoid wasp, *Trichomalopsis sarcophagae*, which represents the most common lineage identified by the blast searches. The remaining 25 best matches are distributed mostly among the genomes of other insects, although other non-insect lineages, including bacteria, are represented as well (Table S8), and they mainly correspond to protein-coding regions of transposons. Thus, unlike some B chromosomes, whose sequences derived largely from one or a few large regions of essential chromosomes within their resident genomes (*27*–*29*), PSR consists primarily of three complex repeats (70.32%) and other sequences that are absent from the *N. vitripennis* genome and, in some cases, derived from other organisms.

Quantification of the RNAseq data revealed that 25 of the 44 PSR-encoded loci are expressed at moderate to high levels (TPM >= 10) in *Nasonia* testes (Table S8). The three highest-expressed loci were selected for further analysis (Fig. 1D, Table S8). Two of these sequences were previously identified in an earlier transcriptome profiling study as PSR4317 and PSR1539 (*18*), whereas the third highest-expressed gene, PSR-tra, has a predicted ORF with limited homology to part of the wasp’s sex-determining gene, *transformer.* Based on their transcript sequences and chromosomal gene models (Fig. S4), PSR4317 and PSR1539 may contain translational open reading frames (ORFs) of 279 and 242 amino acids, respectively (*18*). Although neither of these two PSR sequences matches any gene present in the *N. vitripennis* genome, PSR4317 shows some homology to a nuclear hormone receptor of several invertebrates and contains several putative functional domains, including a zinc finger C4 type DNA binding domain, RNA polymerase I specific transcription initiation factor RRN3 domain, and a transcription factor IIB zinc-binding domain. Interestingly, the highest homology covering the whole length of PSR4317 corresponds to a gene present in the endosymbiont *Candidatus cardinium*, which is known to produce reproductive manipulations, including cytoplasmic incompatibility (CI), parthenogenesis, and feminization, in certain arthropod host species (*30*). The observation that the majority of expressed transcripts encoded by PSR are most homologous to *T. sarcophagae* genes (Table S7, S8) is consistent with the previously proposed hypothesis that *Nasonia* PSR originated from the *Trichomalopsis* lineage by interspecific hybridization (*31*).

Using RT-qPCR (Real time quantitative PCR), we detected expression of these three genes in the male germ line, as well as in all tested embryonic and adult male somatic tissues, demonstrating that their expression is not regulated to a specific developmental time (Fig. S5). If any of these genes play a role in PSR’s drive, their action on the paternally inherited chromatin could, in principle, occur either during sperm formation or instead in the egg’s cytoplasm, immediately following fertilization. To address these possibilities, we used RNAi to transiently reduce transcript levels for each of these genes (PSR-4317, PSR-1539, PSR-tra) in the male germ line, and assessed for an effect on drive by measuring the sex ratio of F1 progeny (Fig. 2A, 2B). Knock-down of PSR4317 transcripts in the testes yielded a striking effect on F1 sex ratio: 46% of treated males produced broods that ranged between 10% and 90% F1 females (Fig. 2B). In contrast, RNAi targeting of the other two genes resulted in all-male broods similar to those from control (untreated) PSR+ wasps, despite the fact that their transcript levels were effectively reduced to levels comparable to RNAi-targeted PSR4317 transcript levels (Fig. 2A, 2B, Fig. S6). Maternal RNAi (*32*) targeting of PSR4317 in the egg’s cytoplasm had no effect on F1 sex ratio (Fig. S7; see Methods). Thus, the effect of RNAi on sex ratio appears to be limited to the male germ line.

**Fig. 2.**
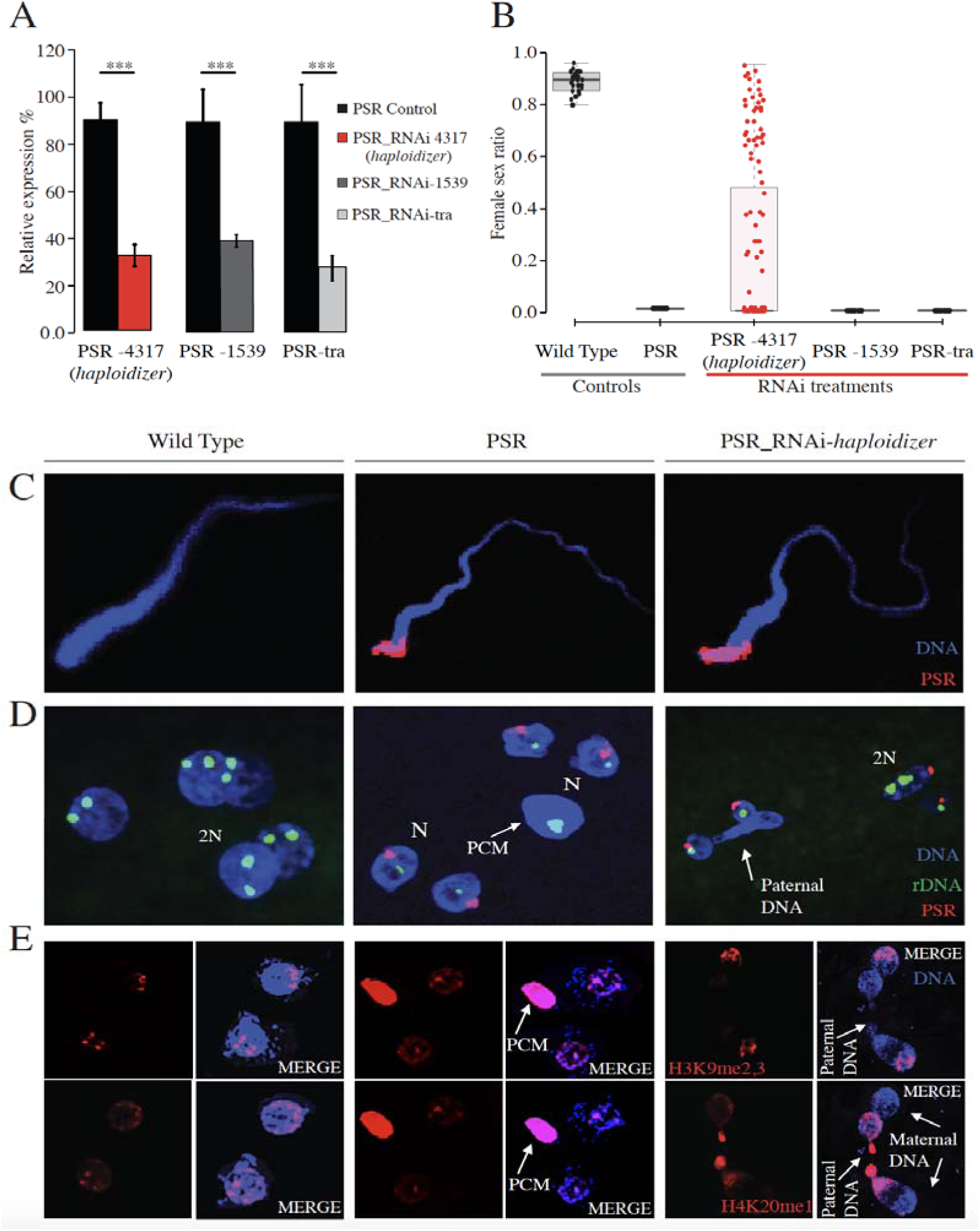
Effects of *haploidizer* targeted by RNAi in early embryos. **A)** Efficiency of RNA interference measured by relative gene expression of the targeted genes. Asterisks indicate significant expression differences between control and RNAi-treated individuals (n=5, *P* < 0.05, one-way ANOVA with Tukey’s multiple-comparisons test). **B)** Female F1 sex ratio was measured for controls and RNAi-treated wasps. The box plots depict the median (thick horizontal line within the box), the 25 and 75 percentiles (box margins) and the 1.5 interquartile range (thin horizontal line). **C)** Confocal imaging of sperm from wild type, PSR+ and *haploidizer* RNAi-treated PSR+ males. **D)** Embryos from wild type, PSR+ and *haploidizer* RNAi-treated PSR+ fathers, stained for with PSR (red) and rDNA (green). **E)** Embryos from wild type, PSR+ and *haploidizer* RNAi-treated PSR+ fathers, stained for the histone marks, H3K9me2,3 (top panels, red) and H4K20me1 (bottom panels, also red). DNA is blue in all panels.. PCM= Paternal chromatin mass, 2N= diploid nuclei, N= Haploid nuclei.

The production of F1 female progeny by RNAi-treated fathers could result from suppression of the genome eliminating activity, but it could also occur from destabilization and loss of PSR before the genome-eliminating activity has occurred. Several observations strongly argue against this latter possibility. First, individual sperm produced by RNAi-treated males contained a single copy of the PSR chromosome (Fig. 2C, Fig. S8), and PSR also was present in the nuclei of young fertilized embryos sired by these males (Fig. 2D, Fig. S9). Additionally, using PCR, we confirmed that the F1 adult females produced in our crosses were positive for multiple different PSR-specific sequences, suggesting that they inherited the B chromosome from their fathers (Fig. S7C, Fig. S10, Table S9). Together, these findings led us to speculate that the appearance of F1 female progeny may result from failure of paternal genome elimination when PSR4317 transcripts are knocked down, thus implicating PSR4317 as an active facilitator of PSR’s drive.

To further test this possibility, we examined the mitotic behavior of the sperm- and egg-derived nuclei and their mitotic descendants in young embryos produced by RNAi-treated fathers. Normally, in embryos from control PSR+ fathers, the sperm-derived nuclear material fails to resolve into individual chromosomes upon entry into the first mitosis, forming a mass of unresolved chromatin that remains distinct from the maternally derived nuclei ((*13*, *14*) Fig. 2D). This paternal chromatin mass (PCM) never segregates, eventually becoming lost within the embryo, as the egg-derived nuclei continue to divide (*13*, *14*). Remarkably, in nearly all of the embryos from PSR+ fathers RNAi-treated for PSR4317, there was no distinct PCM (Fig 2D, Fig. S9). Instead, in about 68% (26 out of 38) of these embryos the paternal chromatin formed bridges that spanned between dividing nuclei (Fig. 2D, Fig. S9). Such bridges were never observed in control PSR+ embryos (Fig. 2D). Additionally, some embryos from RNAi-treated fathers contained multiple nuclei that varied widely in size, suggesting that they were mosaics of nuclei with varying ploidy levels. In support of this idea, we observed one, two, and in some cases more rDNA (ribosomal DNA) foci per nucleus in these embryos (Figure 2D). We also examined two different histone post-translational modifications (PTMs), H3K9me3 (tri-methylation of histone 3 lysine 9) and H4K20me1 (methylation of histone 4 lysine 20), which under control (PSR+) conditions become abnormally distributed across the paternal chromatin in the presence of PSR (*11*). In young embryos from RNAi-treated fathers, however, both of these histone PTMs appeared more similar to patterns that are present in wild type (non-PSR) embryos (Fig. 2E). Taken together, these observations suggest that RNAi-targeting of PSR4317 alleviates the defective state of the sperm-derived chromatin caused by PSR, thereby allowing the sperm-derived chromatin to partially segregate with the egg-derived chromatin, giving rise to embryos that are mosaics of nuclei with differing amounts hereditary material derived from the two parents. Thus, we named this gene *haploidizer* because its expression is required for conversion of diploid embryos into haploids by causing paternal genome elimination.

We further speculated that the PSR+ females generated by RNAi-treated, PSR+ fathers arise from a portion of the mosaic embryos in which genome elimination is partially suppressed. Consistent with this hypothesis, using RT-PCR we were able to amplify both female- and male-specific transcript isoforms of the sex-determining genes, *transformer* (*tra*) and *doublesex* (*dsx*), in all tested PSR+ F1 females, but only male-specific isoforms in control PSR+ and wild type males (Fig. S10). In *N. vitripennis*, initiation of female sex determination is regulated epigenetically (*33*) and requires an unknown trans-acting factor termed *womanizer* (*wom*). This factor promotes female specific splicing of *transformer* (*traF*), whose presence, in turn, directs female-specific splicing of *doublesex* (*dsxF*), thereby promoting female development (*33*). It has been hypothesized that *wom* is epigenetically silenced during oogenesis, and its absence in unfertilized eggs ensures male-specific development (*33*). We therefore propose that the PSR-induced alteration of the sperm-derived chromatin in control (PSR+) embryos is sufficient to block the production of *dsxF*, thereby facilitating male development. However, in fertilized embryos from RNAi-treated, PSR+ fathers, a less defective state of the paternal chromatin may allow some expression of *wom* and subsequently of *traF* and *dsxF* (Fig. 3). We note that F1 PSR+ females were unable to transmit PSR to their progeny, unlike their male siblings (Fig. 3). It is unlikely that the mosaic nature of these PSR+ females is the cause of this transmission failure because these individuals produce viable offspring, thus indicating normal transmission of the five essential chromosomes. Instead, this effect may reflect an intrinsic block to PSR transmission through the female germ line.

**Fig. 3.**
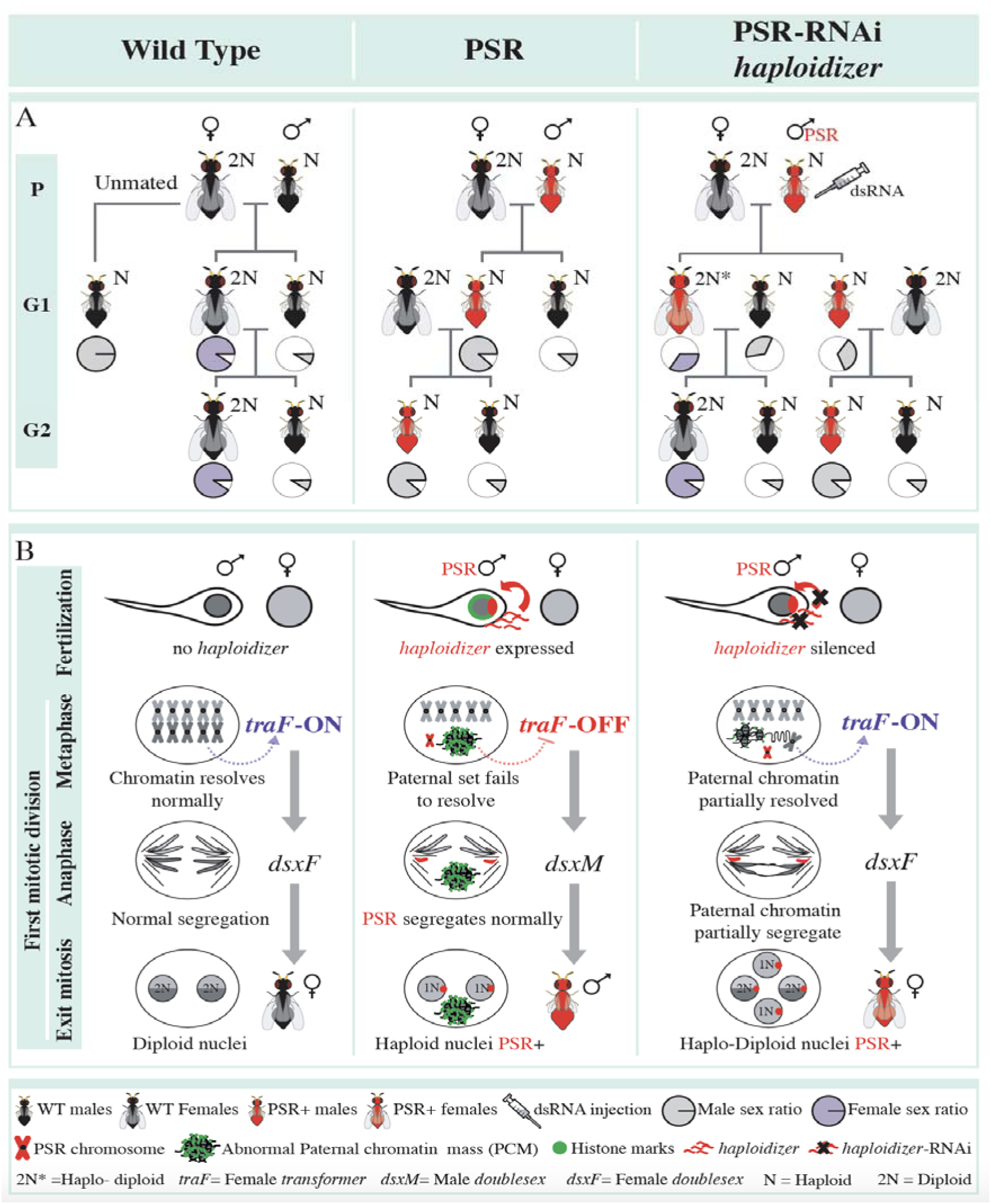
Genetics and active model of PSR-induced genome elimination. **A)** The effects of RNAi knockdown of haploidizer on PSR transmission and sex ratio. The pie charts indicate male (grey) and female (purple) sex ratios **B)** Model for involvement of *haploidizer* in paternal genome elimination. Expression of PSR’s *haploidizer* gene leads to failure of the paternal chromatin to resolve into chromosomes. This action inhibits the activation of female *transformer* (*traF*) and, thus, female development. As a result, these fertilized eggs develop into haploid male offspring that carry PSR. RNAi treatment of *haploidizer* alleviates the chromatin defects, allowing partial expression of *traF* from the paternal set. This action re-establishes the female developmental pathway.

Although it is not yet known how *haploidizer*’s expression in the male germ line leads to alteration and eventual elimination of the sperm-derived chromatin, our study suggests an intriguing possibility: that *haploidizer* may be derived from a symbiont’s gene that causes cytoplasmic incompatibility (CI) in its insect host. While speculative, this hypothesis is consistent with the fact that both symbiont-induced CI (*30*) and PSR’s effect similarly involve disruption of the paternally inherited chromatin, and they occur at the same mitotic division.

## Acknowledgments

We thank Vijaya Kumar for help with library preparations. Sequencing was performed at the Millard and Muriel Jacobs Genetics and Genomics Laboratory at the California Institute of Technology.

## Author contributions

O.S.A. and P.M.F. conceptualized the study. E.D.B., I.A., T.Y. and Q.M.N. performed the various genetics, molecular experiments and bioinformatic analysis in the study. All authors contributed to the writing, analyzed the data, and approved the final manuscript.

## Data and materials availability

All Illumina, Pacbio and Oxford Nanopore sequencing data can be found at NCBI under accession number PRJNA575073 (https://dataview.ncbi.nlm.nih.gov/object/PRJNA575073?reviewer=uslhbl0iop93gab77lb31ts0n5). Previously published RNA-seq data (18) were also used in this study. They can be found under the accession number PRJNA212516 (https://www.ncbi.nlm.nih.gov/bioproject/PRJNA212516). The genome assembly submission ID is SUB6362372. All other data is available in the main text or the supplementary materials.

## Supplementary Materials

Materials and Methods

Figures S1-S11

Tables S1-S10

Data S1-S2

References (*34-42*)

