## Supplemental Files for "Genome Elimination Mediated by Gene Expression from a Selfish Chromosome"

**This PDF file includes:**

Materials and Methods

Figs. S1 to S11

Caption for Table S1 to S10

Caption for Data S1 to S2

**Other Supplementary Materials for this manuscript include the following**:

Table S1 to S10

Table S1. BUSCO scores

Table S2. PSR-specific contigs identification

Table S3. Assembly statistics

Table S4. Placed contigs

Table S5. Summary of PSR composition

Table S6. Repeats family and abundance on PSR

Table S7. PSR specific transcripts

Table S8. PSR specific genes

Table S9. Sex Ratio of G1 females and males after RNAi

Table S10. Primers used and their application

Data S1 to S2

Data S1. Genome assembly files

Data S2. Gene prediction

**Materials and Methods**

**Sequencing and Assembly of the *N. vitripennis* Genome**

Genomic DNA from 50 *Nasonia vitripennis* individuals from AsymCx strain, either with or without PSR, was isolated as a total using a Blood & Cell Culture DNA Midi Kit (Qiagen, Cat# 13343) according to the manufacturer’s protocol. DNA integrity was assessed using Genomic DNA ScreenTape assay (part number 5067-5365) for 4200 TapeStation System (Agilent, Inc., Santa Clara, CA), quantified with Qubit dsDNA HS Kit (ThermoFisher Scientific #Q32854) and used for PacBio® whole genome sequencing. Genomic library was sequenced on PacBio RSII instrument using 30 P6-C4 flowcells generating 3.47 million reads with total yield of 30.98 GB. Genomic library for nanopore sequencing was prepared using 1D Genomic DNA by Ligation library construction kit (SQK-LSK109) and sequenced on 2 R9.4 flowcells using MinION desktop sequencer (Oxford Nanopore Technologies Ltd, Oxford, UK) generating 1.41 million reads with total yield of 11.66GB (Table S3). Basecalling was performed with Albacore v2.3.1. Illumina libraries were constructed from wild type and PSR-carrying samples using NEBNext Ultra™ II DNA Library Prep Kit (NEB #E7645) and sequenced on Illumina HiSeq2500 in paired end mode with the read length of 250 nt to the sequencing depth of 30 million reads per sample. PacBio Reads were assembled with canu v1.6 (*34*) and polished with Quiver (Pacific Biosciences, Inc.). PacBio assembly was further scaffolded with nanopore reads using npScarf v1.7-05b (*35*) and polished with Pilon v1.22 (*36*) using Illumina sequencing reads. PSR-specific contigs were identified by mapping Illumina reads generated from WT and PSR+ genomic DNA and calculating chromosome quotient, which is a normalized ratio of the number of WT and PSR+ reads mapping to a contig (*22*) (Table S2, Fig. S1). The assembly comprises 444 contigs with N50 of 6.60 MB and total length of 297.31 MB. 120 of those contigs are PSR-specific with N50 of 124 kb and total length of 9.2 MB (Table S3).

**Genetic map placement**

Genetic markers developed based on differential hybridization of species-specific oligos between *N. vitripennis* and *N. giraulti* (*23*) were used to place contigs on the genetic map. 73 contigs totaling 272,286,400 bp (91.58%) were assigned to five linkage groups. Of those, 13 (159,467,447 bp, 53.64%) were properly oriented, while 60 (112,818,953 bp, 37.95%) were located in pericentromeric regions without recombination between markers and were placed without orientation or precise relative position (Fig. S2, Data S1.tar). Genome completeness was assessed using BUSCO pipeline (Table S1) (*21*). Repetitive elements were discovered and masked by running RepeatModeler and RepeatMasker (*24*, *25*).

**Genome annotation with nanopore RNA-seq**

PSR+ testes and whole animal RNA-seq libraries for nanopore sequencing were constructed with cDNA-PCR Sequencing Kit (SQK-PCS109) and sequenced on two R9.4 flow cells for 48 hours each using MinION desktop sequencer (Oxford Nanopore Technologies Ltd, Oxford, UK). Basecalling was performed with Guppy v3.0.3 in high accuracy mode. The libraries generated 16.05 and 11.43 million reads with yields of 12.16 and 12.01 GB, respectively (Table S4). Illumina RNA-seq libraries were constructed using NEBNext Ultra II RNA Library Prep Kit for Illumina (NEB #E7770) and sequenced on Illumina HiSeq2500 in paired end mode with the read length of 100 nt to the sequencing depth of 30 million reads each. Full length cDNA reads were identified with Pychopper (Oxford Nanopore Technologies) and aligned to the genome with minimap2 (*37*). Pinfish pipeline (Oxford Nanopore Technologies) was used to generate gene models supported by at least 10 full length aligned nanopore reads. The pinfish pipeline generated 18593 transcripts corresponding to 11482 genes (Data S2). 68 transcripts (44 genes) are encoded by PSR contigs. Nanopore reads were remapped to pinfish transcripts with minimap2 and transcript expression levels were quantified using salmon (*38*). Illumina RNA-seq data was quantified against pinfish transcripts with featureCounts (*39*).

**Expression of PSR candidate genes in early development, carcass and testes.**

Twenty females from AsymC strain were individually mated to a PSR male and give one *Sarcophaga bullata* pupa every other day for 4 days. Thereafter these females were allowed to oviposit for 1h at 25°C. Five replicates of 30 embryos per time point were collected in Trizol reagent (Invitrogen, USA) and stored at -20°C until extraction. For time points 2h and 15h, embryos were incubated respectively 1h and 14h before collection to reach the right developmental time. Testis and carcasses were dissected from one day adult PSR males. Five replicates with 3 pairs of testes each and five replicates with 3 PSR carcasses each were collected in Trizol reagent and stored at -20°C until extraction. Total RNA was extracted from each sample with Trizol reagent according to the manufacturer’s instructions. Each sample was subjected to DNase treatment to eliminate any DNA contamination, and approximately 1ug of total RNA was reverse-transcribed with oligo-dT and hexamer primers in a 1:6 ratio with the RevertAid^TM^H Minus First Strand cDNA Synthesis Kit (Fermentas, Hanover, MD, USA). The cDNA was then diluted 30x before being used for real time PCR (qPCR). qPCR was performed with SYBR Green (Genesee Scientific). 4ul of diluted cDNA was used for each reaction of 20ul total containing primers at a final concentration of 0.2μM and 10μl of SYBR Green buffer solution. Three technical replicates for each reaction were performed to correct for experimental errors. *Elongation factor 1α* (*ef1α*) and *adenylate kinase 3* (*ak3*) were used as reference genes for normalization of the data, after confirmation that their expression level is constant throughout development and between tissues (Fig. S11) (*40*). Reactions were run on a Roche LightCycler® System with the following qPCR profile: 120 sec of activation phase at 95°C, 40 cycles of 15 s at 95°C, 30 s at 62°C. The primers are listed in Supplementary Table S9. Relative expression level of the three PSR genes to the reference genes was calculated by normalizing the expression data with LightCycler® 96 software. Gene expression between samples was then compared with two-way ANOVA and Tukey HDR for multiple comparison test in R statistical software (R Development Core Team 2012).

**Parental RNAi of PSR candidate genes**

RNAi knockdown of the three candidate genes was induced in early PSR male pupae by injecting dsRNA of candidate PSR genes. A fragment of 660bp, 400bp and 601bp was produced respectively for each gene (PSR4317, PSR1539 and PSRtra). At either 5’ and 3’ end of the fragment a T7 promoter was placed using designed primer in supplementary table S10. The fragments were transcribed in both directions using the Megascript RNAi kit (Ambion, Austin, Texas, USA). Briefly, sense and antisense RNA fragments were synthesized in separate transcription reactions. After 6h incubation at 37°C, the two reactions were mixed and heated at 75°C for 5 min followed by cooling down slowly (overnight). Exonuclease digestion removed DNA and ssRNA and dsRNA was subsequently purified according to the kit protocol. Finally, dsRNAs were precipitated with ethanol and re-dissolved in water and stored at -20°C.

PSR male pupae were injected in the abdomen following the procedure of (*32*), either with 4 μg/μl of dsRNA_4317 (RNAi_4317), dsRNA_1539 (RNAi_1539) or dsRNA_tra (RNAi_tra) mixed with red dye. Injections were performed under continuous injection flow with Femtojet Express system (Eppendorf) using aluminosilicate glass filaments pulled with Sutter Instrument. Pupae were injected at the posterior end until the abdomen turned clearly pink. Slides with injected wasp pupae were incubated in an Agar/PBS Petridish at 25^o^C. Control pupae were injected with red dye mixed with water in a 1:4 ratio. After emergence, males were mated singularly with one WT female for 24h and afterward stored in -80^o^C for further processing.

**Genetic crosses**

Females mated with RNAi-treated males were hosted with two *S. bullata* pupae and allowed to oviposit for two days. After oviposition females were used to produce embryos for FISH experiment (See below) and hosts were incubated at 25^o^C. Ten day later, the hosts were opened and *Nasonia* pupae were scored for the presence of females (G1 females). DNA extraction from 35 G1 females, offspring of RNAi-treated males, were tested for the presence of PSR with PCR using PSR-specific primers (Fig. S5A).

To test if PSR in G1 females and males is functional and normally inherited, about 35 G1 females and 34 males PSR^+^ were singularly cross respectively with wild type males and females. G2 offspring sex ratio was scored 10 days after oviposition (Table S9). PSR presence was assessed for all crossed individuals in each generation with PCR using PSR-specific primers (Table S10).

**Expression analysis of PSR genes after RNAi**

In order to assess the efficiency of the RNAi reaction, total RNA was extracted from control and RNAi injected males (one male per sample) and cDNA conversion was performed as described above. Expression data were first analysed as described above using *Ef1α* and *ak3*as reference genes. Relative expression level of the three PSR genes to the reference genes was calculated by normalizing the expression data with LightCycler® 96 software. Gene expression between samples was then compared with two-way ANOVA and Turkey HDR for multiple comparison test in R statistical software (R Development Core Team 2012).

**Embryo collection and fixation**

Wild type mated females, crossed with RNAi treated- males PSR+, were placed into individual glass vials and allowed to oviposit into a single blowfly pupa for discrete lengths of time. Specifically, a 45 min-1 hr laying time was used for obtaining embryos stages between fertilization and immediately before the first mitotic division, whereas a laying time of 1 hr 15 min was used for collecting embryos undergoing the first mitotic division or in the second S-phase. Embryos were carefully removed from host pupae by using one half of a separated pair of ultrafine forceps and placed into a 10 mL screw top glass vial. The following solutions were quickly placed into the vial with embryos in the following order: 3 mL heptane; 1.5 mL 1x Phosphate Buffered Saline (1x PBS); 600 μL 37% formaldehyde. The vial with embryos in fixative was placed onto a platform rocker and embryos were fixed for exactly 28 minutes. Following this time, embryos were removed and placed onto a small piece of Whatman paper and allowed to dry for ~30 sec to 1 min. The embryos were lightly pressed downward onto double-sided adhesive tape secured to the surface of a clean 22 mm Petri plate. 1.5 mL of 1x PBT (1x PBS with 0.1%Triton-X 100) was then placed into the Petri plate to keep the embryos hydrated. The embryos were carefully de-vitellinized under a dissecting micro- scope by using a 28-gauge hypodermic needle. The de-vitellinized embryos were transferred to a 0.6 mL microfuge tube, washed three times with 1x PBT and stored at 4–6 °C before staining.

**DNA *in situ* hybridization (FISH)**

A small, single stranded DNA probe used to detect PSR in fixed embryos was based on the following sequence that is exclusive to the PSR chromosome: 5′–CAC TGA AAA CCA GAG CAG CAG TTG AGA–3′. The telomere was detected by using a ssDNA probe with the following sequence: 5’-TTA TTG GGT TAT TGG GTT ATT GGG TTA TTG-3’. The rDNA locus was marked with a cocktail of the following two ssDNA probes, each recognizing a different part of the 18S IGS repeat: 5’- TTA GAC TTT TTC GAG CCT CCG AGA-3’ and 5’-ATT GAC GCT CGC ACA TCA CTC ATT-3’. These sequences were chemically synthesized by IDT Inc. (USA) and fluorescently labeled at their 5′ ends with Alexa-488, Cy3, or Cy5. Before DNA FISH, immuno-stained embryos were post-fixed in the dark for 45 min in 4% paraformaldehyde and then washed 3 times in 2x saline-sodium citrate and Tween-20 (SSCT). From this point, whole mount DNA FISH was conducted exactly as previously described (*41*). For DNA FISH of squashed chromosomes, we followed exactly a previously described protocol, using testis as a source of mitotic chromosomes (*42*).

**Immunostaining**

To visualize H3K9me3 and H4K20me1 we stained fixed embryos with primary antibodies (rabbit anti-H3K9me3 and mouse anti-H4K20me1, Active Motif, Inc.) diluted 1/500 in PBT, overnight on a platform rocker at 4 °C. Subsequently, embryos were then washed 3 times at 10 minutes each with 1x PBT, and then stained with fluorescently conjugated secondary antibodies at room temperature for 1 hr on a platform rocker in the dark. Secondary antibodies used in this study were: anti-rabbit Cy3 and anti-mouse Cy5 (both at 1:300; Invitrogen-ThermoFisher, Inc., USA). Embryos were then washed as stated above and then mounted on a slide with Vectashield mounting medium containing DAPI (Vector Laboratories, Inc., USA)

**Confocal microscopy and image processing.**

Fluorescence microscopic imaging was conducted with a Leica TCS SPE confocal microscope. Images were collected as Z-series for each laser channel and subsequently merged for visual capture of cellular features within the same nucleus that were not in the same focal plane. Merged images were exported in highest quality JPEG format, and then re-merged and processed in Adobe Photoshop CS5 v. 12.

**Supplementary figures:**

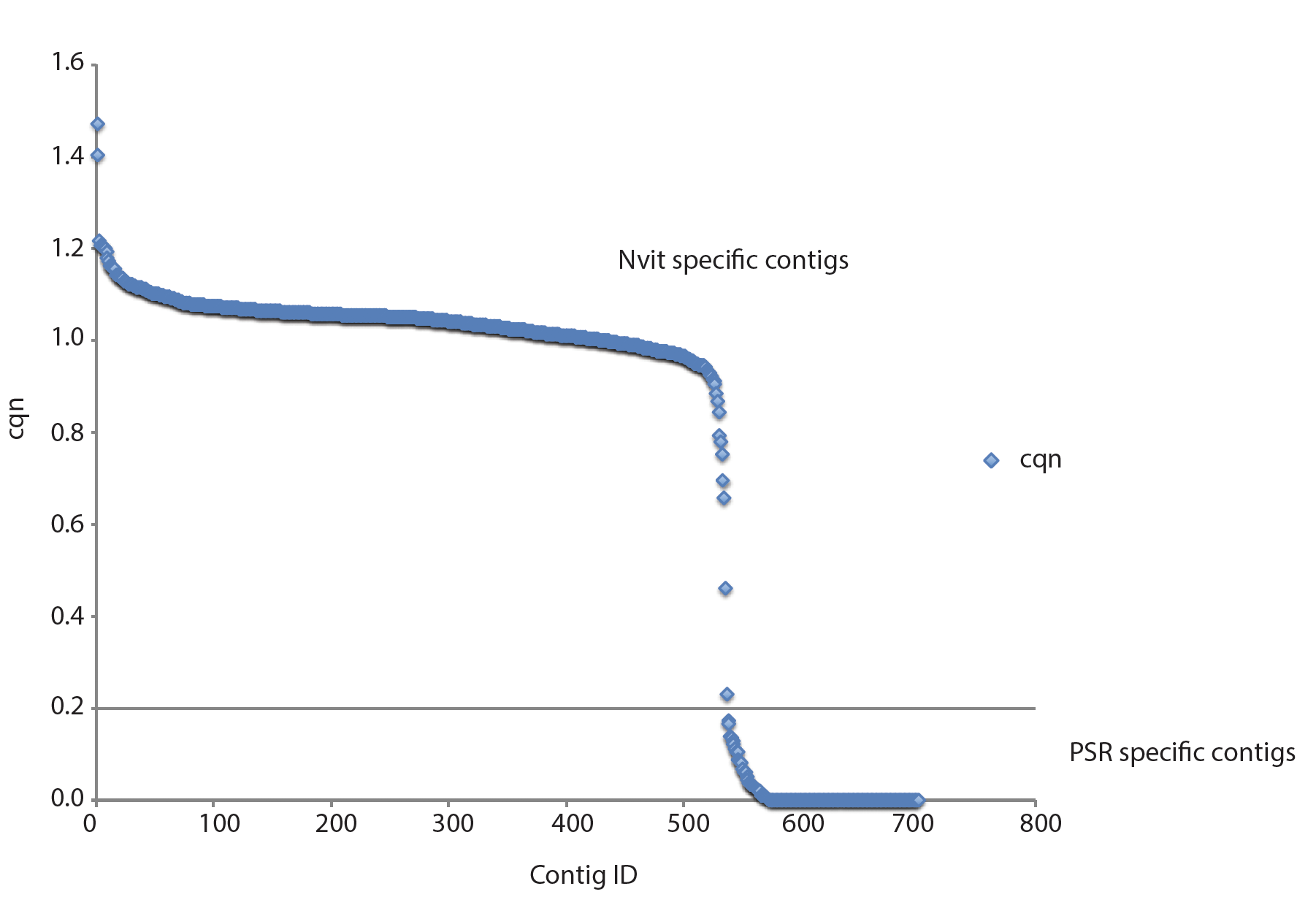

**Fig. S1. Chromosome quotient of contigs**

Chromosome quotient (cqn) represents a normalized ratio of the number of wild type and PSR+ reads that map to each contig. PSR-specific contigs are the ones with cqn smaller than 0.2.

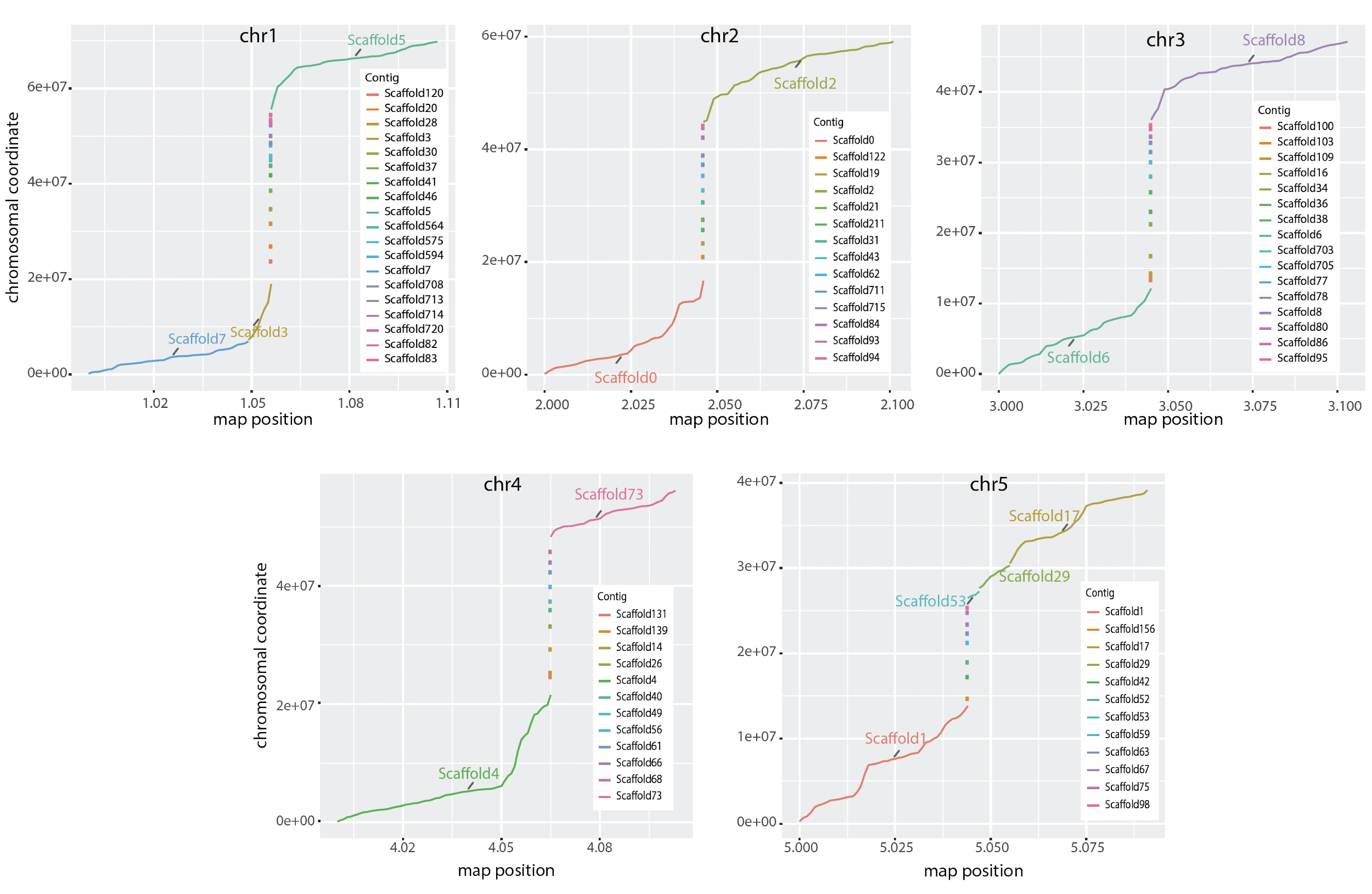

**Fig. S2. Genetic map of *Nasonia vitripennis***

*N. vitripennis* contigs totaling 272 Mb (91.58% of total genome) were assigned to the five essential *N. vitripennis* chromosomes using the previously developed genetic markers (*22*).

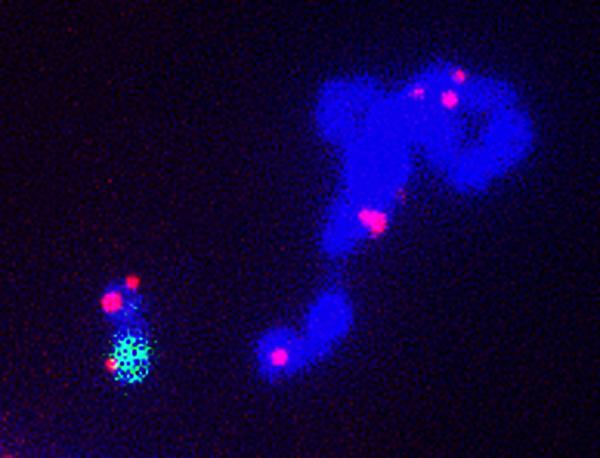

**Fig. S3. Telomeres**

Confocal imaging of Telomere sequences (TTATTGGG) on the 5 A-chromosome of *N. vitripennis* and on PSR chromosome. DNA is highlighted by DAPI (blue) and telomeres in red.

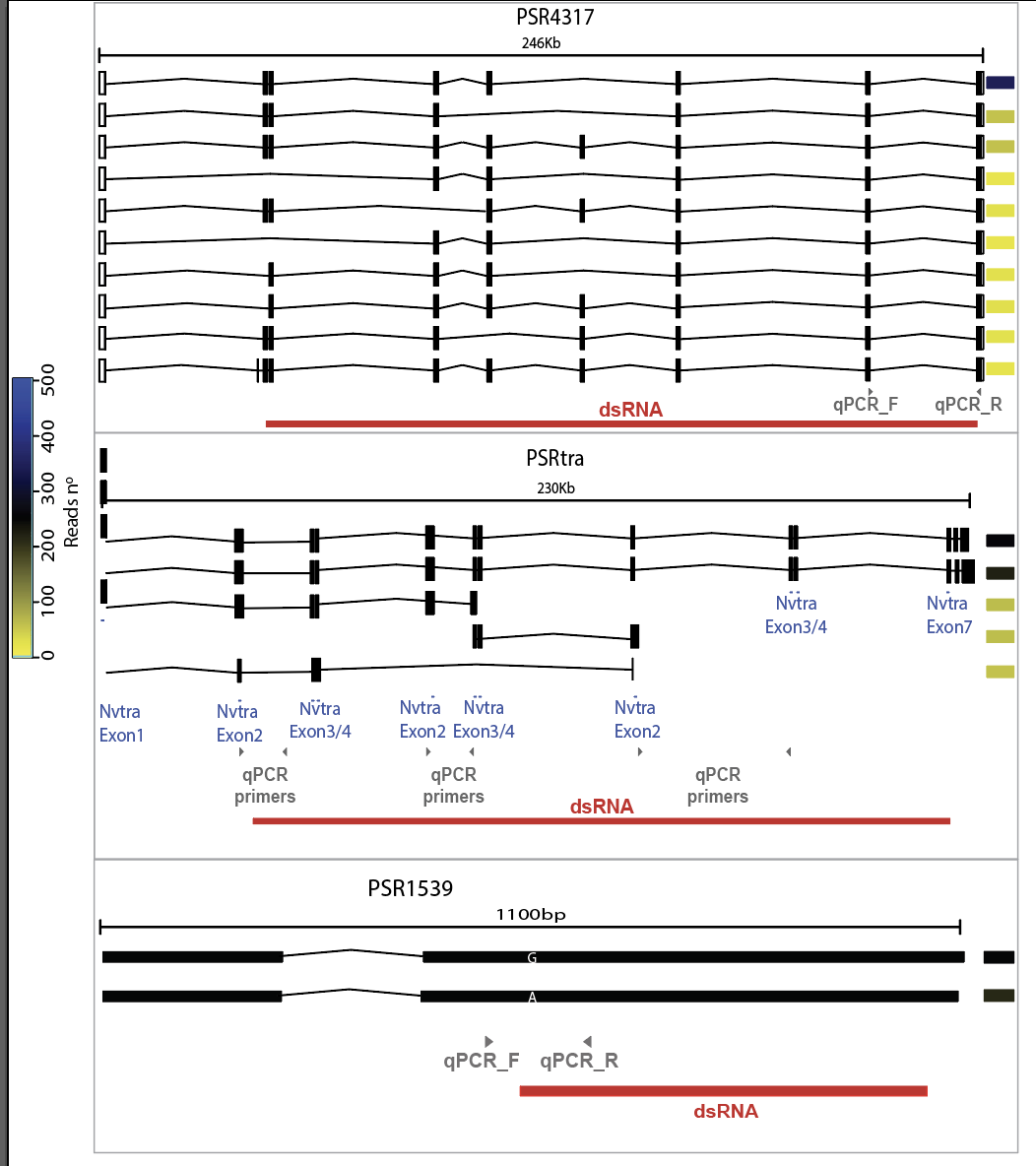

**Fig. S4. gene model of candidate PSR genes**

Schematic representation of five PSR candidate genes. Exons are indicated with boxes and intron with lines. Each row represents a splice variant and colored boxes on the right of each scheme indicate expression level in reads number (Yellow to blue for low to high expression value). The total length is indicated above each scheme in KB. Red lines indicate the region targeted by RNAi. Arrows indicates the qPCR primers location.

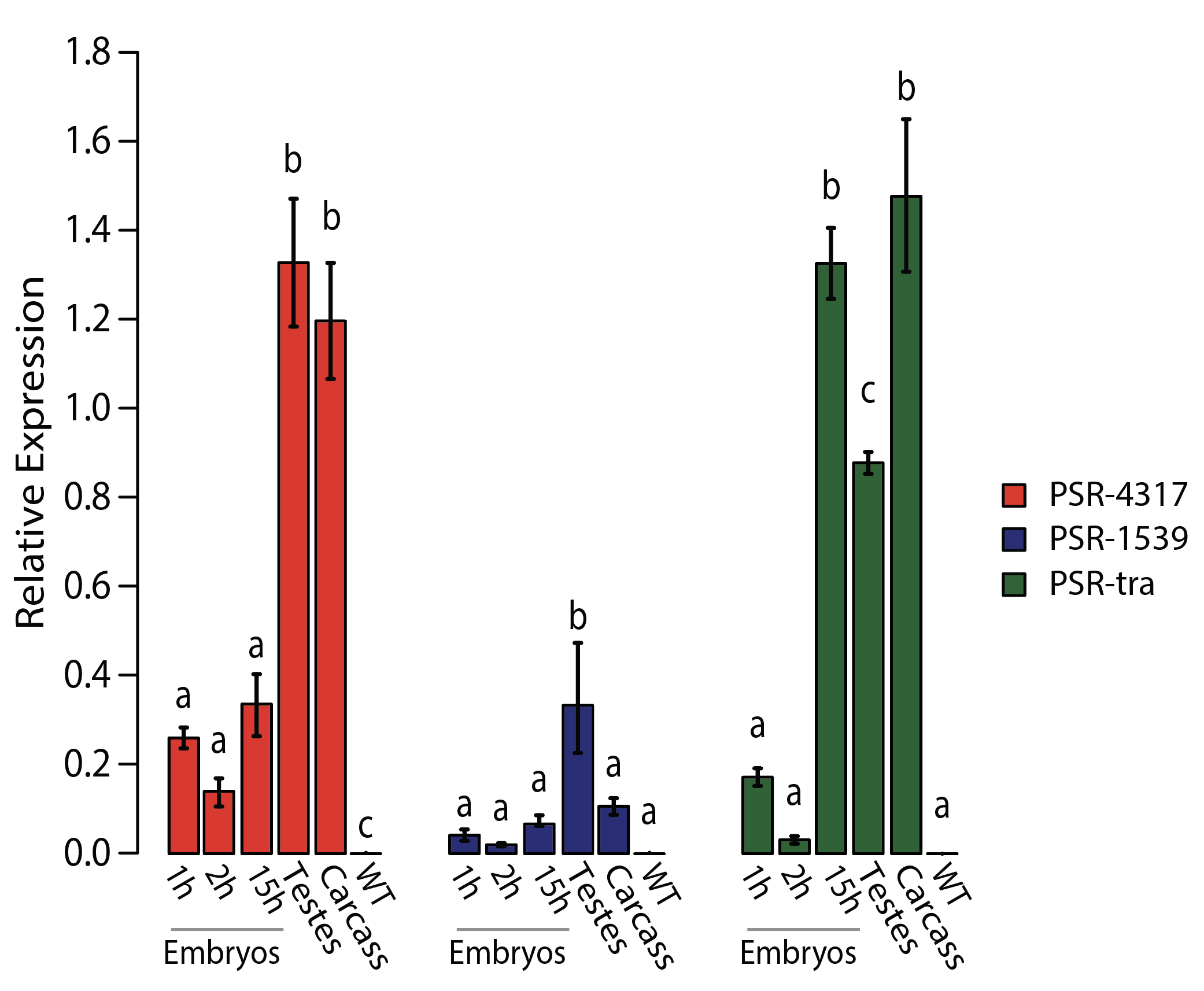

**Fig. S5. Validation of expression of PSR candidate genes**

Relative expression from RT-qPCR of PSR candidate genes among three embryonic time points, testes and carcass samples. Different letters indicate significance differences per tested gene between time points (n=5, *P* < 0.05, one-way ANOVA with Tukey’s multiple-comparisons test).

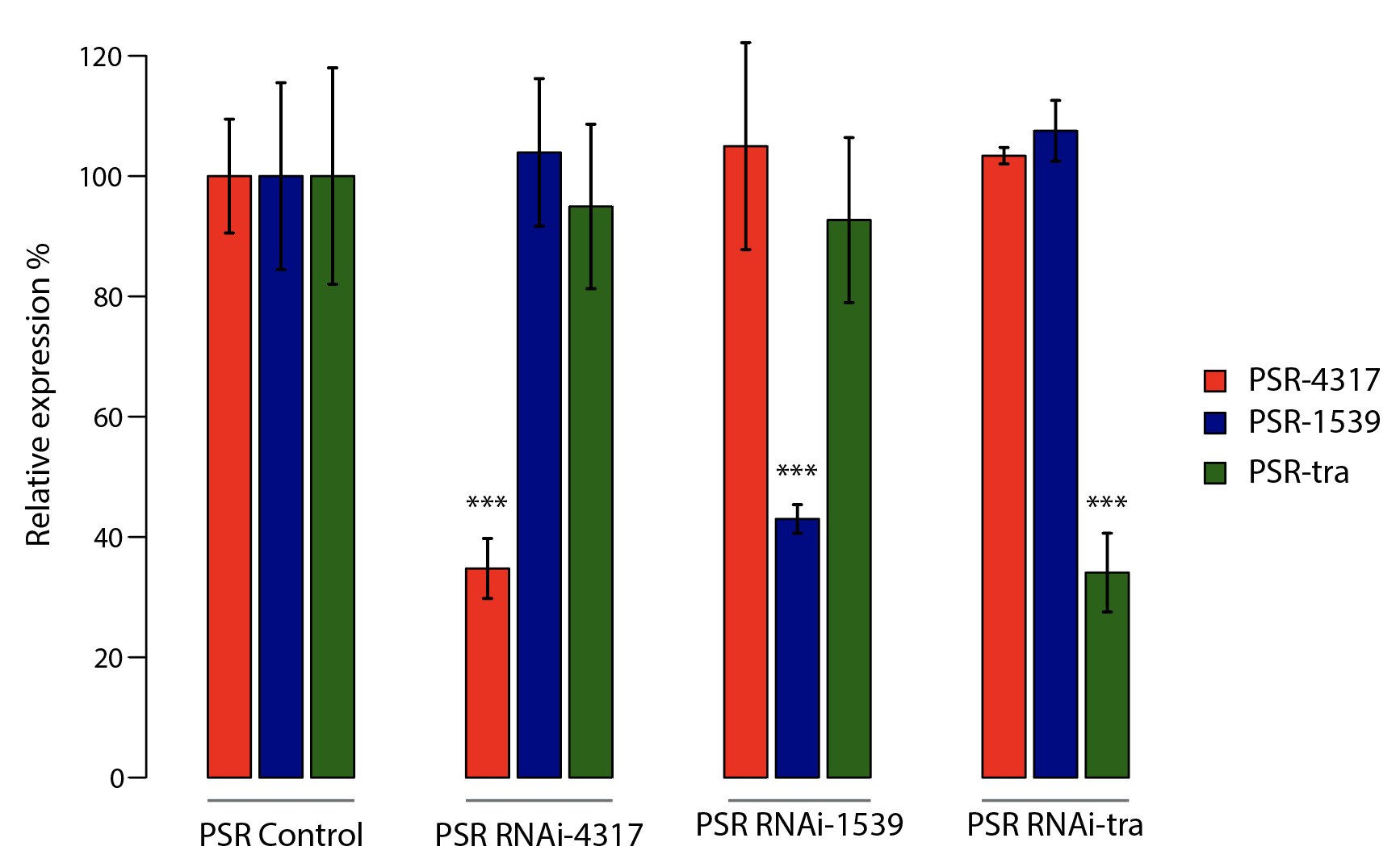

**Fig. S6. RNAi efficiency**

Relative gene expression of PSR candidate genes in untreated (PSR Control) and RNAi treated PSR males (PSR RNAi-4317, PSR RNAi-1539, PSR RNAi-tra) 24h after emergence. Different colours indicate different gene tested per each RNAi experiment. Asterisks indicate significant difference (n=5, *P* < 0.05, one-way ANOVA with Tukey’s multiple-comparisons test).

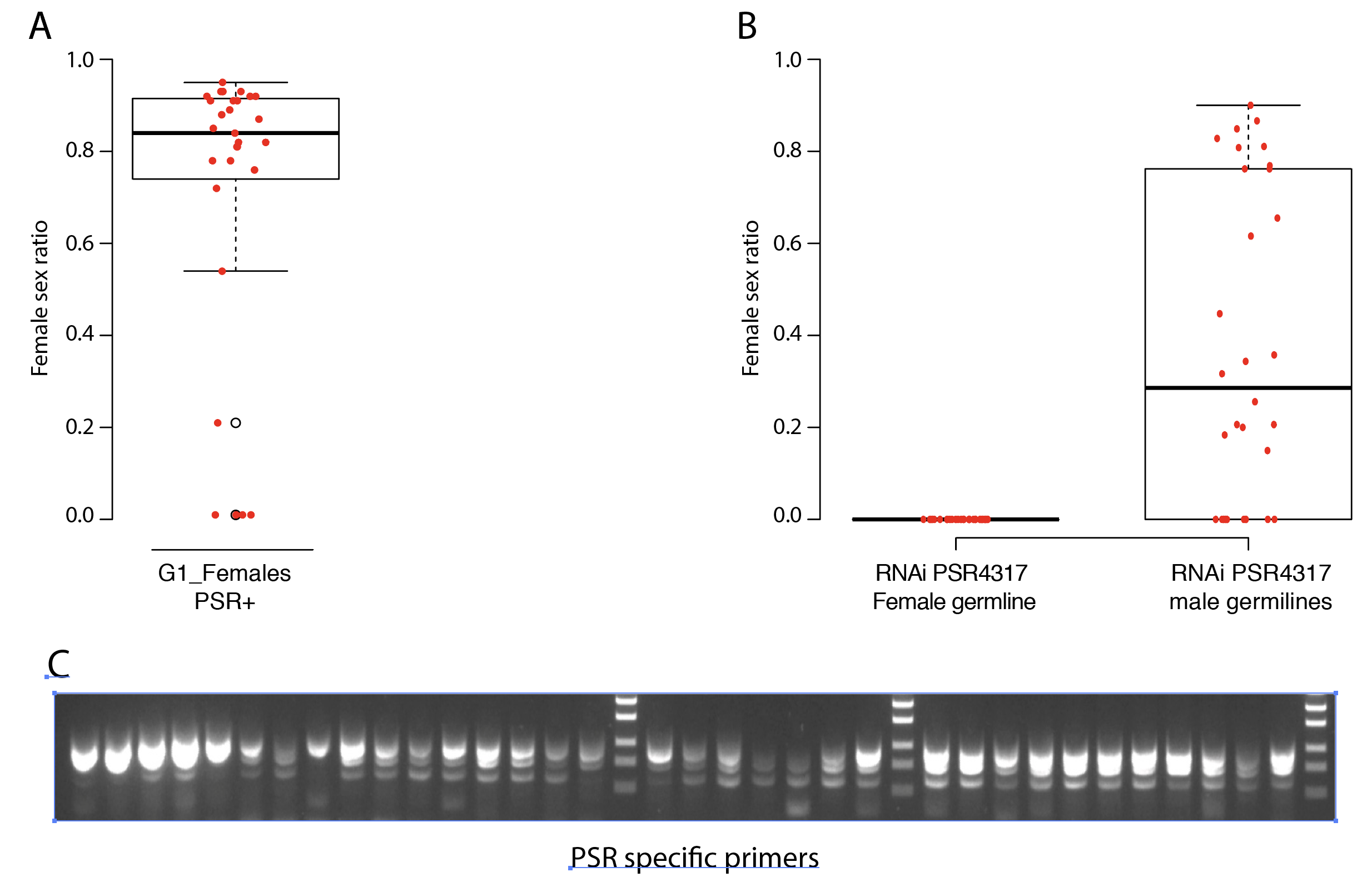

**Fig. S7. Sex Ratio after PSR-RNAi**

**A)** Box plot displaying female sex ratio produced from PSR+ G1 females originate from positive RNAi males. **B)** Box plot displaying female sex ratio after RNAi either in male or female germlines. **C)** Gel electrophoresis showing that tested G1 females carry PSR using PSR specific primers.

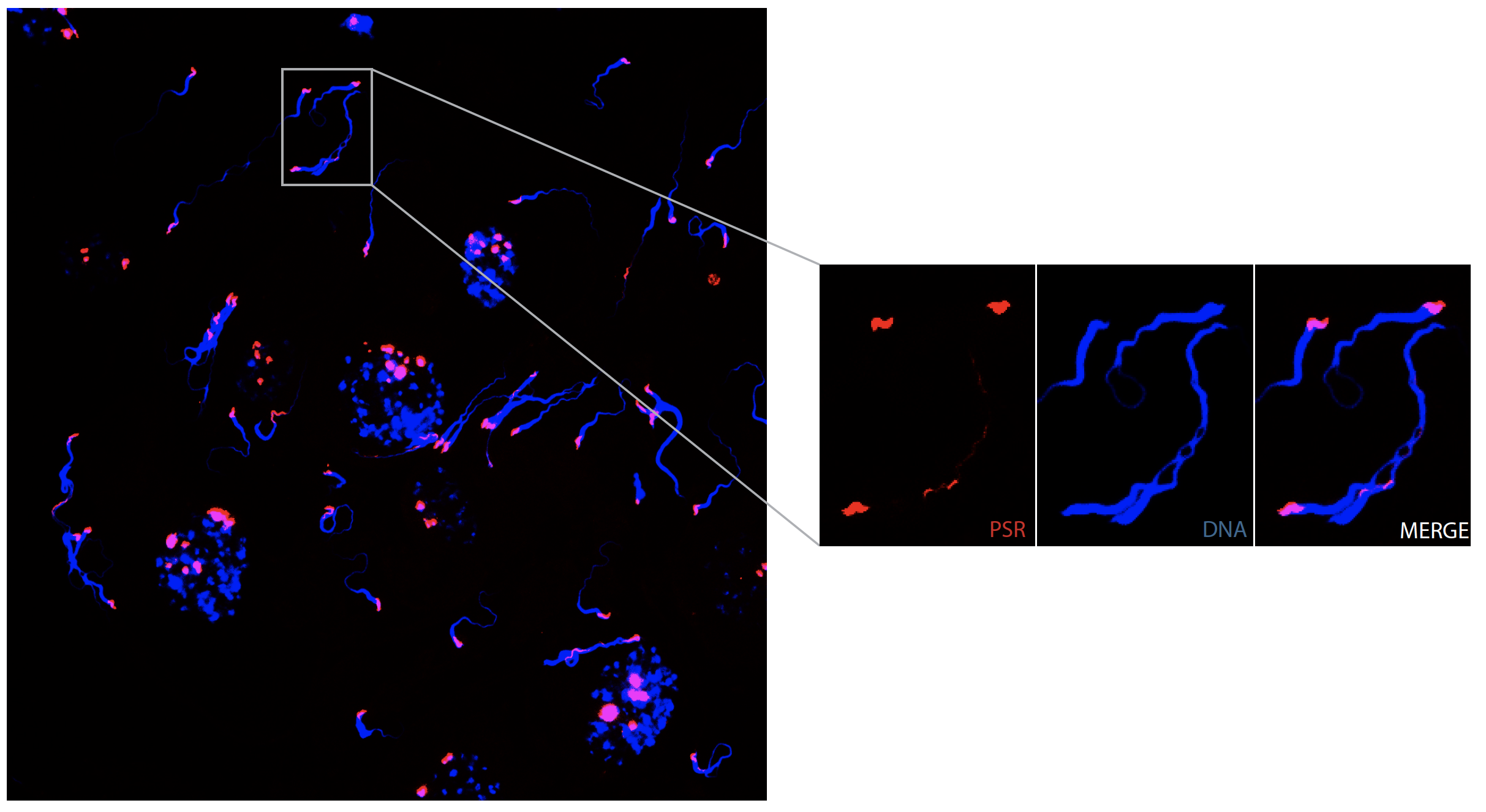

**Fig. S8: Sperm fluorescent *in situ* hybridization**

Fluorescent *in situ* hybridization of PSR+ RNAi-treated sperms. DNA is highlighted by DAPI (blue) and PSR by a sequence-specific FISH probe (red).

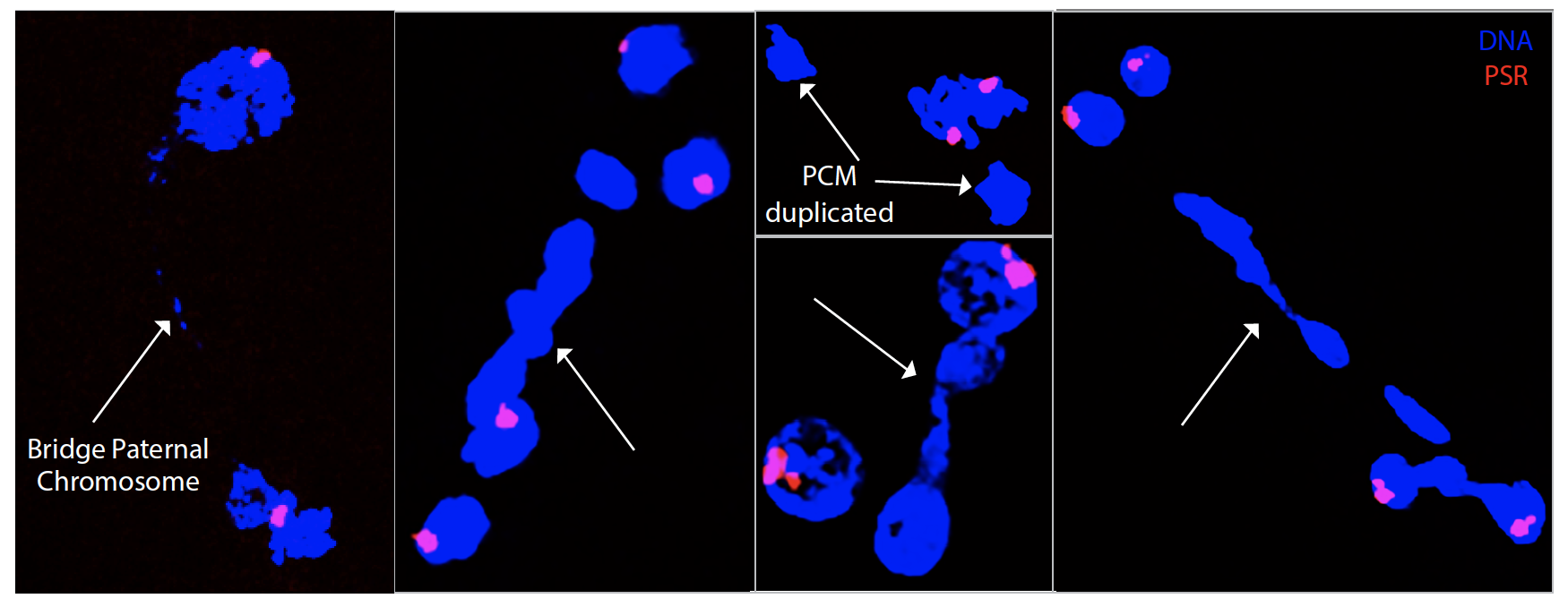

**Fig. S9. Embryonic fluorescent *in situ* hybridization of RNAi-treated PSR+**

Additional examples of Fluorescent *in situ* hybridization of RNAi treated PSR+ males. DNA is highlighted by DAPI (blue) and PSR by a sequence-specific FISH probe (red). White arrows indicated disrupted PCM (Paternal chromatin mass).

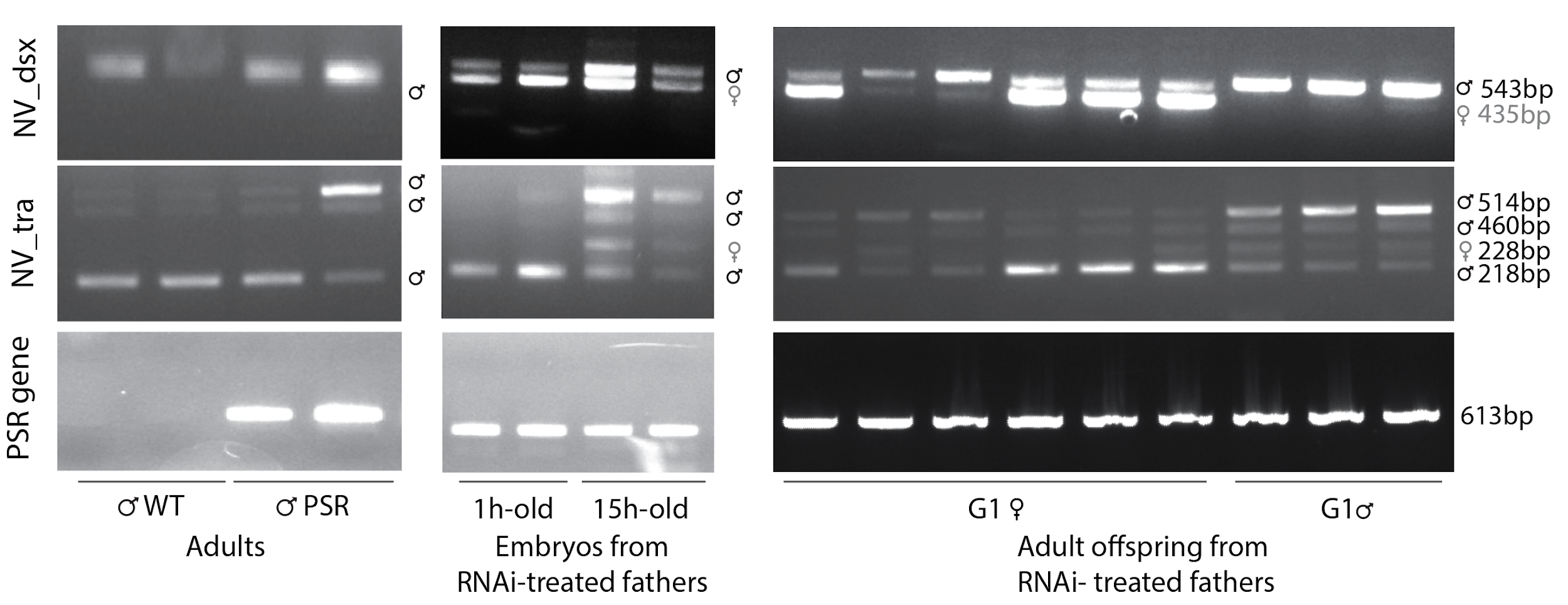

**Fig.S10. *dsx* and *tra* splicing**

Gel electrophoresis from RT-PCR depicting differential splicing variants of *NVdsx* (*doublesex*), *NVtra* (*transformer*), and PSR specific gene in wild type males (WT), PSR males, 1h and 15h-old embryos from RNAi-treated fathers and G1 females and males originated from RNAi-treated fathers. *NVdsx* and *NVtra* diagnostic primers from *(27)*. PSR gene analysed using PSR specific primers (Table S10).

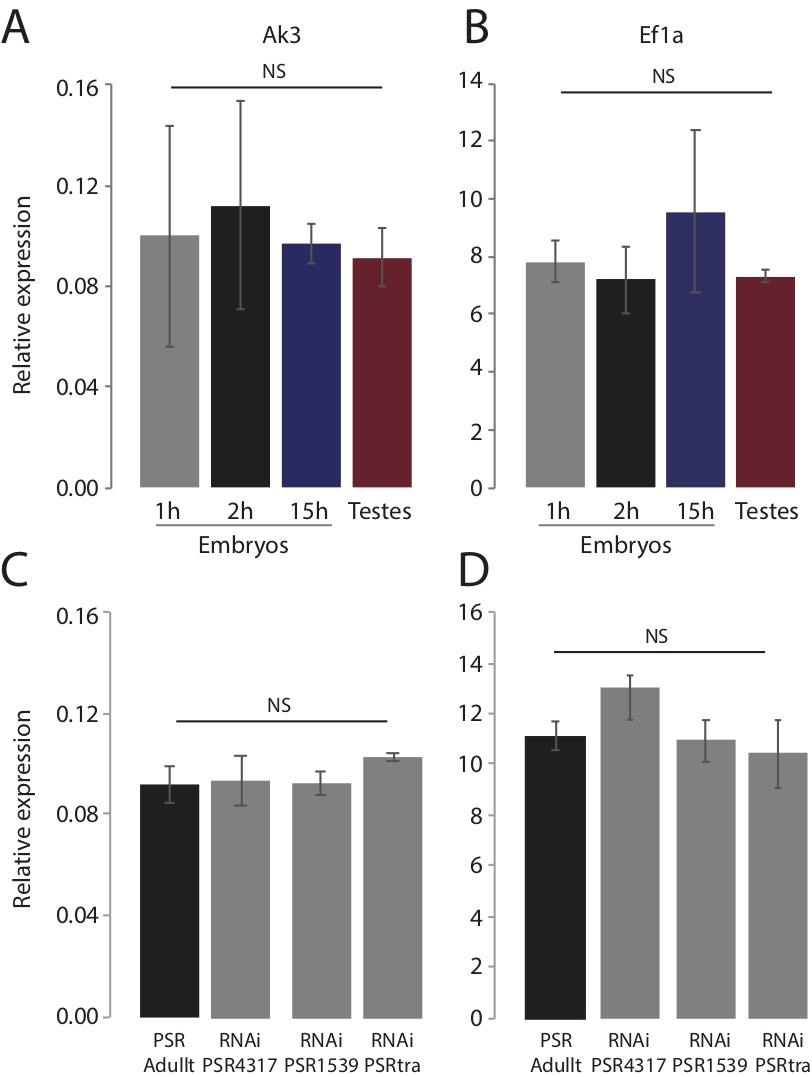

**Fig.S11. Stability of reference genes between samples and treatments**

The average relative expression of **A)** Ak3 and **B)** EF1α is compared among time points and **C)**, **D)** among RNAi- treated samples by two-way ANOVA.

**Additional File**

**Supplementary Tables**

Table S1 to S10

**Table S1. BUSCO scores**

Genome completeness assessed using BUSCO pipeline (*21*).

**Table S2. PSR-specific contigs identification**

List of all contigs and their cq ratio (chromosome quotient (*20*)) used to identify PSR-specific contigs. Cq calculates the ratio of the number of wild type and PSR+ reads mapping to a contig.

**Table S3. Assembly statistics**

Information about assembly of wild type Nasonia vitripennis genome and PSR chromosome

**Table S4. Placed contigs**

Information about contig numbers and length

**Table S5. Summary of PSR composition**

Summary of sequences present on PSR and their abundance.

**Table S6. Repeats family and abundance on PSR**

List of PSR repeats family, location and type of repeats.

**Table S7. PSR specific transcripts**

List of PSR expressed transcripts from Nanopore RNA sequencing of PSR-carrying testes and whole animal.

**Table S8. PSR specific genes**

List of PSR genes with location on PSR scaffold, Blast results, Pfam domain and expression data (TPM). TPM expression value are calculated with featureCounts (*27*) using Nanopore RNAseq data and an additional illumina RNA seq dataset from (*18*).

**Table S9. Sex Ratio of G1 females and males after RNAi**

Sex ratio of individual G1 females and males and presence of PSR

**Table S10. Primers used and application**

List of primers used for PCR, qPCR and dsRNA production.

Data S1-S2

**Data S1. Genome assembly files**

The archive contains a file with all the contigs in the assembly (Nvit_psr_1.fsa), the PSR specific contigs (Nvit_psr_1.psr_specific.fsa), the positions of placed contigs within chromosomes (Nvit_psr_1.chromosomes.agp) and chromosomal scaffolds generated using genetic markers (*23*) (Nvit_psr_1.chromosomes.fsa). All gaps are represented by 100 Ns. Relative position of contigs lacking orientation is unknown.

**Data S2. Gene prediction**

The file includes gene models supported by at least 10 full length aligned nanopore reads

generated with Pinfish pipeline (Oxford Nanopore Technologies).
